# An Inducible Model of RPE-Associated Hydroxyapatite Mineralization with Relevance to Drusen Formation in Age-Related Macular Degeneration

**DOI:** 10.64898/2026.10.08.757467

**Authors:** Julian Yang, Steffi Daniel, John D. Hulleman

## Abstract

A cardinal feature of nonexudative age-related macular degeneration (AMD) is the accumulation of drusen between the retinal pigment epithelium (RPE) and Bruch’s membrane, yet the mechanisms that initiate and promote drusen formation remain to be fully understood. Hydroxyapatite (HAP), a calcium-phosphate mineral identified within human drusen, has been proposed to contribute to drusen nucleation and is associated with increased likelihood of RPE atrophy. However, mechanistic studies providing insight into RPE-associated mineralization have been limited by a lack of easy-to-use and manipulatable culture models. Here, we established a rapid, inducible ARPE-19-based model of HAP mineralization. We found that ARPE-19 cells cultured in minimum essential media (MEM) supplemented with nicotinamide (NIC), sodium phosphate, and ascorbic acid (NaPAA) reproducibly formed HAP positive deposits, whereas mineralization was not observed while using standard DMEM/F12 medium, DMEM/F12 with NaPAA, MEM-NIC medium, or MEM-NIC medium with alternative osteogenic supplements (such as β-glycerophosphate, ascorbic acid, and dexamethasone). Scanning electron microscopy with energy dispersive X ray spectroscopy demonstrated that the formed deposits were enriched in calcium (Ca) and phosphorus (P) with Ca/P ratios comparable to drusen-associated HAP deposits from AMD donor tissue. ARPE-19-based HAP deposits were also closely associated with neutral lipid-rich material, staining positive for BODIPY around their core structure. Bulk RNA sequencing of mineralized ARPE-19 cells revealed extracellular matrix, angiogenic, inflammatory, and metabolic transcriptional changes in the absence of strong enrichment of canonical osteogenic pathways. Co-treatment of mineralizing cells with either pyrophosphate, EDTA, or sodium citrate significantly reduced mineral deposition, but these approaches were not able to reverse it. Together, these findings establish a tractable RPE mineralization model that recapitulates key features of AMD-associated HAP and provides a platform for investigating mechanisms of drusen-associated mineral formation.

## INTRODUCTION

Age-related macular degeneration (AMD) is one of the leading causes of irreversible vision loss in older adults worldwide and is broadly categorized by nonexudative and exudative (angiogenic) forms of the disease(Bikbov et al., 2025; Kong et al., 2025; Korb et al., 2022). The canonical hallmark of nonexudative AMD is the accumulation of drusen, extracellular deposits that form between the basal lamina of the retinal pigment epithelium (RPE) and the inner collagenous layer of Bruch’s membrane (BrM)(Mullins et al., 2000). Although drusen presence, size, number, and morphology are strongly associated with AMD onset and progression(Fleckenstein et al., 2024), the processes that initiate drusen formation and growth remain unclear. The composition of drusen is a complex mixture of lipids(Wang et al., 2010a), proteins(Crabb et al., 2002), and mineral components(Thompson et al., 2015), suggesting that drusen formation involves multiple cross-interacting components(Flores-Bellver et al., 2021; Grillo et al., 2021; Kotnala et al., 2023; Lekwuwa et al., 2022; Thompson et al., 2015). An interesting advance in the characterization of drusen was the identification of hydroxyapatite (HAP) spherules or nodules found in some post-mortem human AMD druse that heralded increased potential for high-risk progression to advanced AMD with RPE atrophy(Fragiotta et al., 2019; Thompson et al., 2015). This was intriguing since HAP is a calcium-phosphate mineral composition normally associated with bone and teeth formation(Blair et al., 2023; Deshpande et al., 2011; Lee et al., 2014; Li et al., 2023; Naim and Sen, 2025; Seredin et al., 2022). HAP in AMD eyes appear as two main calcified structures: spherical deposits approximately 0.5–20 µm in diameter with a cholesterol core, and much larger multilobal nodules(Tan et al., 2018; Thompson et al., 2015). These calcified deposits are proposed to provide a scaffold on which proteins and other drusen components can accumulate, potentially contributing to drusen nucleation and growth(Thompson et al., 2015). Such deposits containing HAP can also be identified non-invasively by optical coherence tomography (OCT) imaging, exhibiting hyperreflective shells followed by hyporeflective centers, sometimes accompanied with bright internal hyperreflective foci(Tan et al., 2018). Together, these observations suggest that HAP may represent a clinically significant component of drusen which is potentially involved in AMD progression.

Despite the identification of HAP in human AMD eyes, how HAP forms in the sub-RPE environment remains to be defined. Studies conducted to characterize the presence of HAP within druse of human AMD eyes have largely focused on its identification and elemental composition rather than the mechanisms underlying its formation. Current hypotheses of HAP-associated drusen formation have proposed an active role for HAP in contributing to drusen formation(Bergen et al., 2019; Thompson et al., 2015). One study by suggested that HAP deposits may arise in association with lipid-rich extracellular material that accumulates on the BrM and subsequently serves as a nucleating surface for additional druse components(Thompson et al., 2015). Notably, these sub-RPE HAP deposits are morphologically and spatially distinct from calcification associated with the elastin layer of BrM observed in diseases such as pseudoxanthoma elasticum, suggesting that the local environment driven by the presence of lipid-rich material may influence where and how mineralization occurs(Tan et al., 2018; Thompson et al., 2015). Experimental studies provided additional evidence that the RPE itself can participate in this process. Primary porcine RPE were shown to generate HAP spontaneously after prolonged culture, while serum-deprived ARPE-19 cells induce amelotin, an enamel-associate protein that promotes HAP formation in teeth(Pilgrim et al., 2017; Rajapakse et al., 2020). However, the conditions that initiate mineralization, the relationship between HAP and other drusen constituents, and the cellular response accompanying mineral formation still remain unexplored. Insights from other soft tissues that ectopically mineralize therefore can provide a useful context for considering the cellular and extracellular factors that may regulate RPE-associated HAP formation. Ectopic calcification has been extensively studied in the contexts of vasculature and cardiac valves, where mineral deposition is now understood as a regulated process involving extracellular matrix composition, extracellular and/or matrix vesicles, local calcium and phosphate availability, and the balance between endogenous promoters and inhibitors of mineralization(Kapustin et al., 2015; Shanahan et al., 2011). In osteoblast and vascular smooth muscle cell models, osteogenic reprogramming can involve runt-related transcription factor 2 (RUNX2) and tissue non-specific alkaline phosphatase/alkaline phosphatase (TNAP/ALPL) induction, while extracellular matrix components such as collagen and/or elastin and vesicles can facilitate calcium-phosphate nucleation and growth(Kapustin et al., 2015; Shanahan et al., 2011). However, ectopic mineralization can also arise through genetic mutations or disturbances in phosphate or pyrophosphate intake/excretion homeostasis, as illustrated by disorders such as primary familial brain calcification and ATP binding cassette subfamily C member 6/ ectonucleotide pyrophosphatase/phosphodiesterase 1 (ABCC6/ENPP1)-associated ectopic mineralization(Albright et al., 2015; Wallingford et al., 2017). These observations from other tissues illustrate that similar calcium-phosphate deposits can arise through distinct cellular programs. However, understanding RPE-associated mineralization requires consideration of tissue-specific differences, as the mechanisms and substrates that support HAP formation can vary even though calcium and phosphate remain its principal mineral constituents. Together, these results highlight the need for RPE-based systems in which the mechanisms governing HAP formation within the sub-RPE environment can be more directly investigated.

Established mineralization models also provide an experimental framework for inducing and studying HAP formation *in vitro*. Approaches have been used to study both osteoblast mineralization and ectopic vascular calcification, particularly in vascular smooth muscle cells, where HAP formation can be induced by modifying extracellular phosphate availability(He et al., 2018; Ho et al., 2023; Montes de Oca et al., 2010; Nagy et al., 2021). In these models, phosphate is commonly supplied as inorganic sodium phosphate or β-glycerophosphate, often together with ascorbic acid to promote extracellular matrix formation and subsequent mineral deposition(Beck et al., 2003; Faure et al., 2025; Ho et al., 2023; Ma et al., 2024; Qiu et al., 2021; Yamazaki et al., 2025; Yevlashevskaya et al., 2023; Zhai et al., 2024). These established osteogenic and vascular calcification models therefore provide a framework for testing whether comparable mineralizing conditions could be adapted to RPE cells. However, similar inducible systems for studying RPE-associated HAP formation have not been established. An experimental model in which RPE mineralization can be reproducibly initiated in a defined timeline and under defined conditions would help facilitate mechanistic investigation into these processes. Existing RPE models such as the primary porcine and serum-deprived ARPE-19 cultures require extended culture periods before HAP formation becomes evident. Therefore, a system by which HAP formation could be better controlled would facilitate mechanistic studies of HAP formation and manipulation. Since ARPE-19 cells are experimentally tractable (i.e., they can be genetically modified, virus-infected, used for high-throughput screening, etc) and can acquire RPE characteristics in nicotinamide-containing medium(Hazim et al., 2019), we asked whether osteogenic supplementation typically used for *in vitro* studies could rapidly and reproducibly induce HAP formation in ARPE-19 cells under controlled culture conditions.

Herein we determined the culture conditions required to promote HAP mineral formation, followed by elemental composition deduction and comparison to calcified AMD druse to determine the extent to which the deposits resembled AMD-associated HAP. We further defined the spatial relationship between neutral lipid accumulation and HAP deposits using fluorescence microscopy and determined cellular transcriptional changes induced under osteogenic conditions. Finally, we identified small molecule treatments can prevent mineralization. Together, our study establishes an experimentally inducible RPE mineralization system as a platform for further investigating the cellular and extracellular mechanisms that contribute to HAP formation in the context of AMD.

## METHODS

### Cell culture

ARPE-19 cells (CRL-2302, ATCC) were cultured at 37°C in 5% CO_2_ in DMEM F/12 (Corning) containing 10% fetal bovine serum (FBS) and 1% penicillin-streptomycin-glutamine (PSQ) unless otherwise noted (e.g. MEM-NIC media, see below)). To prepare cells for experiments, conditioned media was aspirated and washed with Hank’s Balanced Salt Solution (HBSS, Sigma-Aldrich, St. Louis MO) before incubating with 0.25% Trypsin EDTA (Gibco, Waltham MA) at 37°C for 5 min. Trypsin was then neutralized using of DMEM/F12 full media before cell counting and seeding at appropriate densities as described below.

### Mineralization conditions

ARPE-19 cells were seeded at a confluent density of 166,000 cells/cm^2^ in DMEM/F12 full media (10% FBS, 1% PSQ) and incubated overnight to allow attachment. Media was then replaced with either DMEM/F12 full media or MEM-NIC (MEM alpha [Sigma-Aldrich] supplemented with 1% FBS, 1% PSQ, 1% N1 supplement [Sigma-Aldrich], 1% MEM NEAA [Gibco], taurine [0.25 mg/mL], hydrocortisone [20 ng/mL], triiodo-thyronine [0.013 ng/mL], and nicotinaminde [10 mM, all small molecules from Sigma-Aldrich]) in the presence or absence osteogenic supplement (2 mM sodium phosphate [J.T. Baker, Radnor PA] and 50 µg/mL ascorbic acid [Fisher Scientific, Waltham MA], referenced as ‘NaPAA’ here onward). Media changes were conducted every 72-96 h for up to 7 or 14 days, obtaining brightfield and far-red (ex/em 628/688 nm) time course images to monitor visible and autofluorescent (HAP) deposits. Stock solutions of sodium phosphate were kept at RT for supplementation with a new stock being made every month.

### Scanning electron microscopy and energy dispersive x-ray spectroscopy

ARPE-19 cultures containing mineralized deposits were prepared for scanning electron microscopy (SEM) by decellularization(Woodard et al., 2022). Briefly, transwell inserts were washed three times with Dulbecco’s phosphate-buffered saline (DPBS; Gibco) and incubated with 0.5% Triton X-100 and 20 mM ammonium hydroxide for 5 min at 37°C with gentle shaking. Decellularized membranes were subsequently washed three times with ultrapure water and allowed to dry prior to SEM analysis.

Deposit morphology and elemental composition were visualized and analyzed using an Apreo 2S Lo-Vac field-emission scanning electron microscope (Thermo Fisher Scientific) equipped with an Oxford Instruments Ultim Max 100 mm² energy-dispersive X-ray spectroscopy (EDS) detector. Samples were examined under low-vacuum conditions to minimize charging of the non-conductive specimens without the need for conductive coating. Regions containing mineral deposits were identified by SEM and subjected to EDS analysis. Elemental spectra were acquired and analyzed using AZtec software (Oxford Instruments, Concord, MA) to determine the elemental composition of individual deposits, with particular attention to Ca and P. Relative elemental abundance was determined from EDS spectra, and Ca/P atomic ratios were calculated for comparison between ARPE-19-derived mineral deposits and mineralized deposits identified in human AMD donor druse.

### HAP staining with BoneTag680 or OsteoSense680

HAP formation was assessed using either BoneTag680RD (LI-COR, Lincoln NE) or OsteoSense680 (Revvity, Waltham MA). For fixed cell imaging, wells were washed twice with HBSS and fixed with 10% neutral buffered formalin (NBF) for 20 min at RT. Following fixation, wells were washed three times with HBSS and stained with 20 µM of BoneTag680 for 20 min at RT. Wells were then washed twice with HBSS before imaging. For live-cell imaging during the mineralization process, HAP was assessed using OsteoSense680. Cells were washed twice with HBSS and incubated with 240 nM of OsteoSense680 in complete culture medium for 10 min at 37°C. Cells were then washed twice with HBSS and imaged in FluoroBrite DMEM (Gibco) to reduce background noise due to phenol red.

### Alizarin Red staining

Calcification was assessed by Alizarin Red staining (Sigma Aldrich). On day 14, the wells were washed twice with HBSS and the cells were fixed with 10% NBF for 20 min at RT. Following three additional washes of HBSS after fixation, the cells were then stained with 4 mM of Alizarin Red for 20 min at RT. The wells were then washed with HBSS repeatedly until no color was observed in the HBSS between washes, typically four times. For quantification of Alizarin staining, bound dye was extracted by adding 5% cetylpyridinium chloride (CPC, Sigma Aldrich) to each well followed by incubation for 1 h at RT. After incubation, the absorbance of the solution was measured at 560 nm on a GloMax Discover (Promega, Madison WI).

### BODIPY staining

BODIPY 493/503 (Invitrogen, Carlsbad CA) was used to detect neutral lipids in mineralized ARPE-19 cultures. For live-cell BODIPY imaging, cells were washed with HBSS and incubated with BODIPY 493/503 and OsteoSense680 in complete culture medium for 10 min. Cells were then washed with HBSS and incubated with FluoroBrite DMEM prior to imaging using a Celigo Imaging Cytometer (Revvity, Waltham, MA). For fixed-cell staining, ARPE-19 cells were washed with HBSS and fixed in 4% paraformaldehyde for 20 min at RT. Cells were then stained with 3 µM BODIPY 493/503 (Invitrogen) and OsteoSense680 diluted in HBSS for 10 min at RT, washed, and prepared for imaging on a Keyence BZ-X800 (Keyence, Itasca IL).

### Bulk RNA-Seq sample prep

ARPE-19 cells were plated at 166,000 cells/cm^2^ in DMEM/F12 full media. The following day, cells were treated with DMEM/F12 or MEM-NIC in the presence or absence of NaPAA. Media treatment was replenished every 72-96 h. After seven days of treatment, RNA was extracted using an Aurum Total RNA kit (Bio-Rad, Hercules CA) and stored at −80°C. Samples were quantified using a Nanodrop (ThermoFisher Scientific) and fifteen microliters of the samples were sent to Novogene (Sacramento CA). One microgram of RNA was used as input material for library generation. Subsequent data analysis was performed using proprietary NovoMagic software which automatically analyzes samples in a pairwise manner.

## RESULTS

### MEM-NIC supplemented with sodium phosphate and ascorbic acid induces HAP mineralization in ARPE-19 cells

The identification of HAP within large druse of AMD patients was a fascinating discovery(Thompson et al., 2015) with important biological relevance to retinal pigmented epithelium (RPE) cell death (i.e., geographic atrophy, GA) and AMD progression(Tan et al., 2018). To learn more about the process of mineralization and its effects on RPE cells, we sought to determine whether a simple ARPE-19 immortalized RPE culture system could be used to model HAP formation. Under normal culture conditions using either DMEM/F12 standard growth media or RPE-differentiation media, MEM-NIC(Hazim et al., 2019), we observed no evidence of mineralization via brightfield microscopy (Fig 1A, B), Alizarin Red staining (ARS, calcium deposits, Fig 1K, L), or BoneTag680RD staining (HAP deposits, Fig 1O, P) after 14 days of culture. Since it was challenging to visualize the presence of cells under brightfield illumination or in instances of negative staining, we used fluorescent staining for zona occludens 1 (ZO-1) and DAPI to provide evidence of cell borders and nuclei (Fig 1G-J). Next, we tested whether inclusion of osteogenic supplement such as sodium phosphate/ascorbic acid (NaPAA) or β-glycerophosphate/ascorbic acid/dexamethasone (BGP), both of which have been used previously to promote mineralization in osteoblasts(Langenbach and Handschel, 2013; Sordi et al., 2021; Yevlashevskaya et al., 2023) and other in vitro models of ectopic mineralization(Patel et al., 2019), could coax ARPE-19 cells into forming HAP deposits. Surprisingly, BGP supplementation did not promote mineralization in either media condition up to 14 days in culture (Fig 1E, F). Moreover, no mineralization was observed using DMEM/F12 media supplemented with NaPAA (Fig. 1C, M, Q). However, incubation of cells in MEM-NIC media supplemented with NaPAA readily generated visible punctate deposits by brightfield imaging (Fig 1D) which also stained positive for ARS (Fig 1N) and BoneTag680RD (Fig 1R). MEM-NIC media in combination with seeding cells at confluence promotes ARPE-19 differentiation(Hazim et al., 2019). Thus, we next asked whether media supplements contained in MEM-NIC media or seeding at sub-confluency were requirements for mineralization. ARS staining showed that removal of MEM-NIC supplements (Sup Fig 1A–E) or seeding ARPE-19 cells at sub-confluent densities in MEM-NIC (Sup Fig 1F– H) did not markedly reduce mineral deposition, suggesting that RPE differentiation was not necessarily required for HAP formation. In addition, to determine whether HAP formation was cell-dependent rather than arising from the media alone, MEM-NIC supplemented with NaPAA was incubated in the presence or absence of ARPE-19 cells for 14 days. OsteoSense680 staining revealed HAP-positive deposits only in cultures containing ARPE-19 cells (Sup Fig 1I), whereas no detectable OsteoSense680 signal was observed in no cell conditions (Sup Fig 1J). These findings suggest that an initially confluent monolayer is also not required for mineralization. Together, these findings indicate that ARPE-19 cells (regardless of density) cultured in MEM-NIC + NaPAA provide a conducive environment for readily forming mineralized HAP deposits (Fig 1D, N, R).

**Figure 1.**
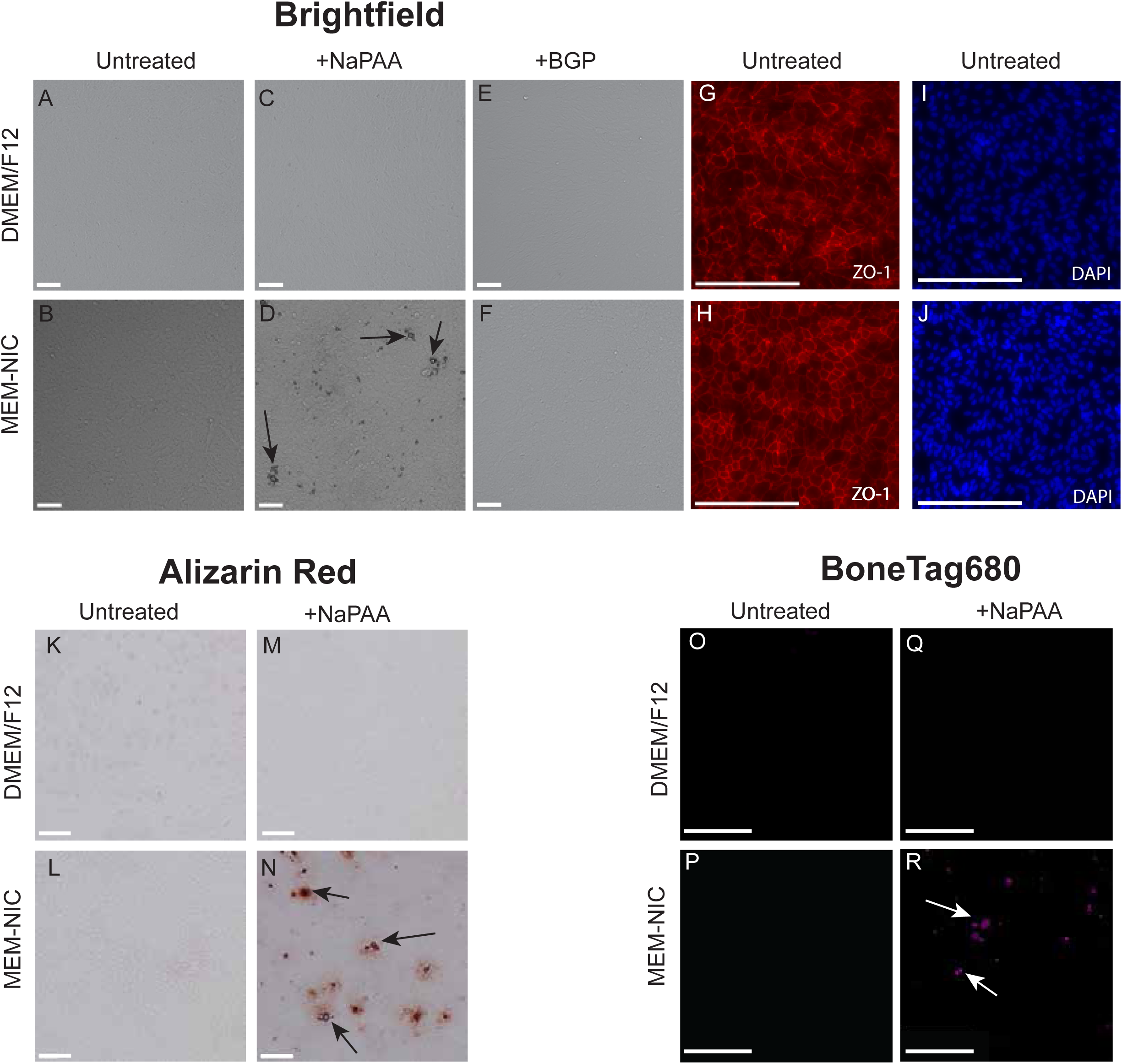
ARPE-19 cells form HAP mineral deposits in MEM-NIC but not DMEM/F12 media under osteogenic supplementation. ARPE-19 cells were cultured in DMEM/F12 or MEM-NIC in the absence or presence of sodium-phosphate + ascorbic acid (NaPAA) or β-glycerophosphate + ascorbic acid + dexamethasone (BGP) for 14 days. (A-F) Representative brightfield images showing visible mineral deposit formation only in MEM-NIC + NaPAA cultures (black arrows). (G-J) ZO-1 and DAPI staining confirmed tight junction formation and the presence of ARPE-19 cells under both DMEM/F12 and MEM-NIC culture conditions and allowed visualization of the epithelial monolayer. (K-N) Alizarin Red staining showed calcium-containing mineral foci only in MEM-NIC + NaPAA cultures. (O-R) BoneTag680 staining further confirmed HAP-associated mineral deposition signal in MEM-NIC + NaPAA cultures. Scale bar = 100 µm, Representative images of n = 3 independent biological experiments.

To further characterize the mineralizing ARPE-19 phenotype, we asked whether HAP formation was accompanied by changes in mitochondrial oxidative phosphorylation (OXPHOS), as phosphate-induced mineralization in ectopic calcification models such as vascular smooth muscle cells has been associated with altered mitochondrial function and ATP production(Alesutan et al., 2020; Shi et al., 2022; Thi Nguyen et al., 2023). NaPAA supplementation did not significantly change mitochondrial OXPHOS in MEM-NIC cells despite robust HAP formation, whereas basal and ATP-dependent OCR increased in non-mineralizing DMEM/F12 cultures supplemented with NaPAA (Sup Fig 2A-E). Thus, these OXPHOS changes did not correlate with the appearance of cell mineralization.

### ARPE-19-derived deposits exhibit calcium and phosphorus enrichment comparable to human AMD-associated deposits

After establishing that ARPE-19 cells cultured in MEM-NIC + NaPAA formed ARS-and BoneTag680-positive mineral deposits, we next sought to confirm the mineral composition of these deposits and determine whether these deposits shared elemental features with HAP deposits observed in human AMD tissue. ARS and BoneTag680 staining approaches do not provide elemental composition or allow direct calculation of the calcium-to-phosphorus (Ca/P) ratio of identified deposits, which is routinely used for assessing the presence of HAP. To remedy this, we used scanning electron microscopy with energy-dispersive X-ray spectroscopy (SEM-EDS) to assess whether ARPE-19-derived deposits were enriched in Ca and P at an elemental level and to determine their compositional similarities or differences to HAP associated with human AMD.

SEM-EDS imaging of ARPE-19-derived deposits demonstrated discrete mineralized structures: spherules approximately 15-20 μm in diameter (Fig 2A-C), and large nodule-like accumulations (Fig 2D-F) with overlapping Ca and P signal in two independent ARPE-19 mineralization samples, an observation consistent with HAP-like mineralization (Fig 2A-F). EDS spectra confirmed that ARPE-19 derived deposits contained strong Ca and P peaks (Fig 2J, K). Importantly, imaging of AMD-derived intermediate drusen deposits confirmed large mineral structures, and elemental mapping showed strong enrichment of Ca and P within the deposit area (Fig 2G-I). EDS spectra also confirmed that human AMD-derived deposits were consistent with HAP-like mineral composition (Fig 2H,I). Additional elements, including sodium, magnesium, aluminum, and chloride, were also detected in all samples, likely a result from sample preparation or residual saline (Fig 2J-L). Ca/P ratio quantification showed that ARPE-19 derived deposits were comparable to human AMD-derived deposits, with ratios of 1.52 for ARPE-19 sample 1, 1.50 for ARPE-19 sample 2, and 1.73 for human AMD (Fig 2M). Together, these findings support MEM-NIC + NaPAA as a relevant inducible culture condition for generating ARPE-19-derived HAP-like mineral deposits that parallel AMD-associated HAP.

**Figure 2.**
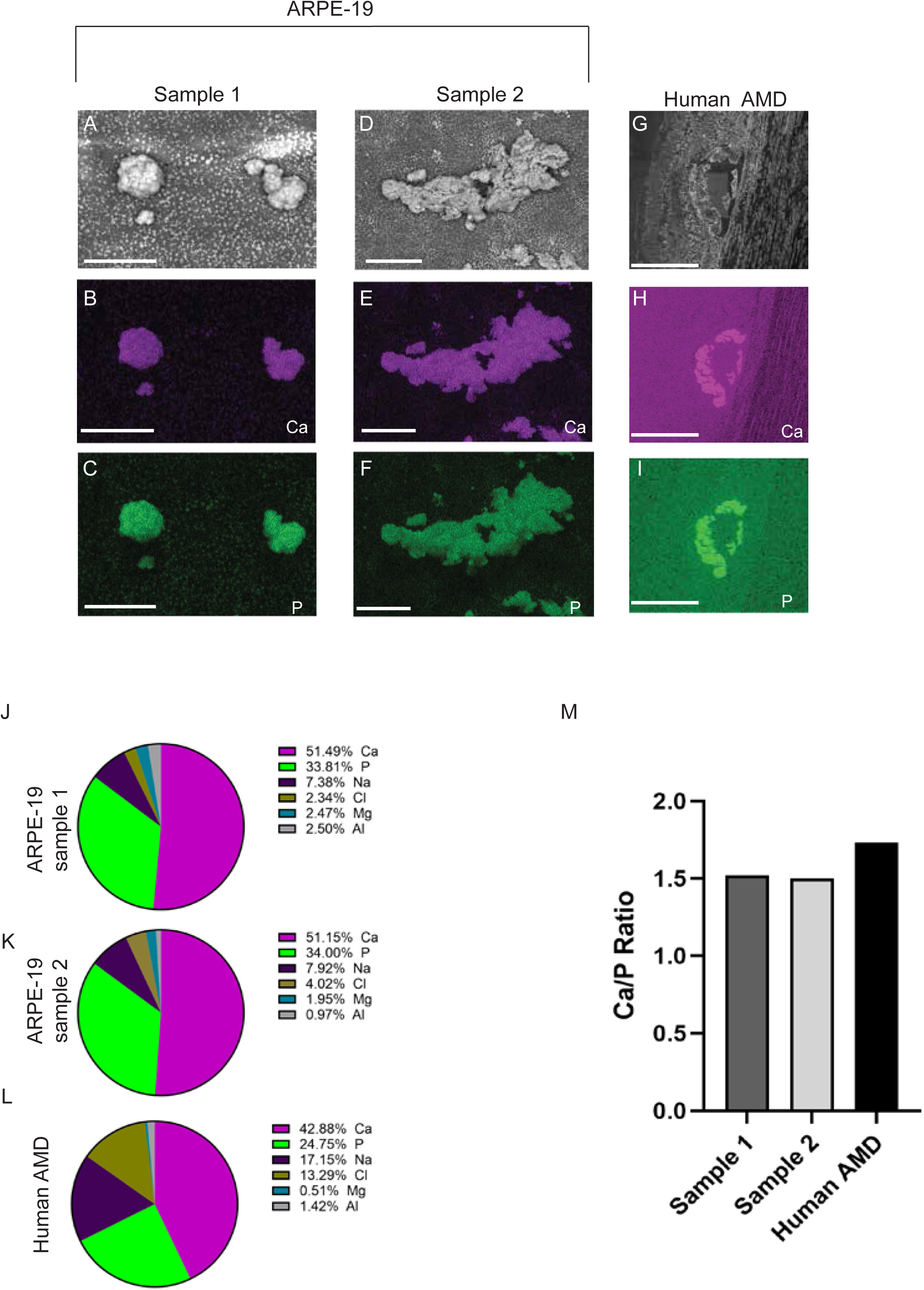
Scanning electron microscopy with energy-dispersive X-ray spectroscopy (SEM-EDS) confirms calcium and phosphorus enrichment in ARPE-19-derived deposits and human AMD drusen. ARPE-19 cells were grown in MEM-NIC + NaPAA for 1 week on transwells. Transwells were decellularlized and processed for SEM-EDS. (A-F) Elemental mapping showed an enrichment of calcium and phosphorus in ARPE-19 derived depsoits consistent with HAP mineral material. Similar trends of calcium (B, E) and phosphorus (C, F) were observed between the two samples. (G-I) Paraffin sections from a human AMD donor (Male, 87 yo, intermediate AMD) containing druse were analyzed by brightfield imaging (G) and SEM-EDS (H-I). Calcium (H) and phosphorus (I) were found to be enriched in druse. (J-M) EDS quantification further showed that calcium and phosphorus were the predominant detected elements in both human AMD drusen and ARPE-19-derived deposits, with comparable Ca/P ratios in human AMD tissue (1.73) and ARPE-19 samples (1.52 to 1.50). Scale bars: A-C = 25 µm; D-F = 50 µm, G-I = 250 µm. n = 2 independent ARPE-19 mineralization samples.

### ARPE-19 derived HAP is spatially associated with neutral lipid accumulation

Previous work identifying HAP in AMD drusen proposed a model in which HAP formation may occur in association with age-related lipid droplet accumulation between the RPE and Bruch’s membrane(Thompson et al., 2015). In this model, lipid droplets could potentially act as nucleation scaffolds for HAP formation, allowing mineral growth to further propagate. Although this lipid-associated nucleation mechanism was not directly tested, it raised the possibility that neutral lipid accumulation may be spatially linked to HAP deposition during drusen development. Having established that ARPE-19 cells cultured in MEM-NIC + NaPAA form HAP deposits (Figs 1, 2), we next asked whether ARPE-19-derived HAP deposits were spatially associated with neutral lipid droplets.

As shown previously, ARPE-19 cells treated with MEM-NIC + NaPAA developed numerous visible punctate HAP deposits by brightfield microscopy (black arrows, Fig 3A, C). Live-cell BODIPY fluorescence imaging revealed neutral lipid signal across the culture surface, with prominent enrichment surrounding HAP deposits as identified by BODIPY (Fig 3B, D), suggesting that lipid-rich material accumulates in close proximity to HAP. To further assess the spatial relationship between neutral lipid and HAP deposits with greater resolution, we performed a separate transwell experiment using ARPE-19 cultures treated with MEM-NIC + NaPAA followed by fixation. OsteoSense680-positive HAP deposits showed overlapping BODIPY signal (white arrows), indicating lipid-rich material within the same regions (Fig 3E, F) and the merged image showed strong spatial association between OsteoSense680-positive HAP deposits and BODIPY-positive neutral lipid signal (Fig 3G). OsteoSense680-only controls showed no detectable background autofluorescence signal in the GFP imaging channel (ex 470/40 nm, em 525/50 nm) used for BODIPY, indicating that the BODIPY-positive signal reflects true lipid labeling rather than autofluorescence or bleed-through from OsteoSense680 (Fig 3 H,I). Together, these findings indicate that ARPE-19-derived HAP deposits are associated with neutral lipid accumulation.

**Figure 3.**
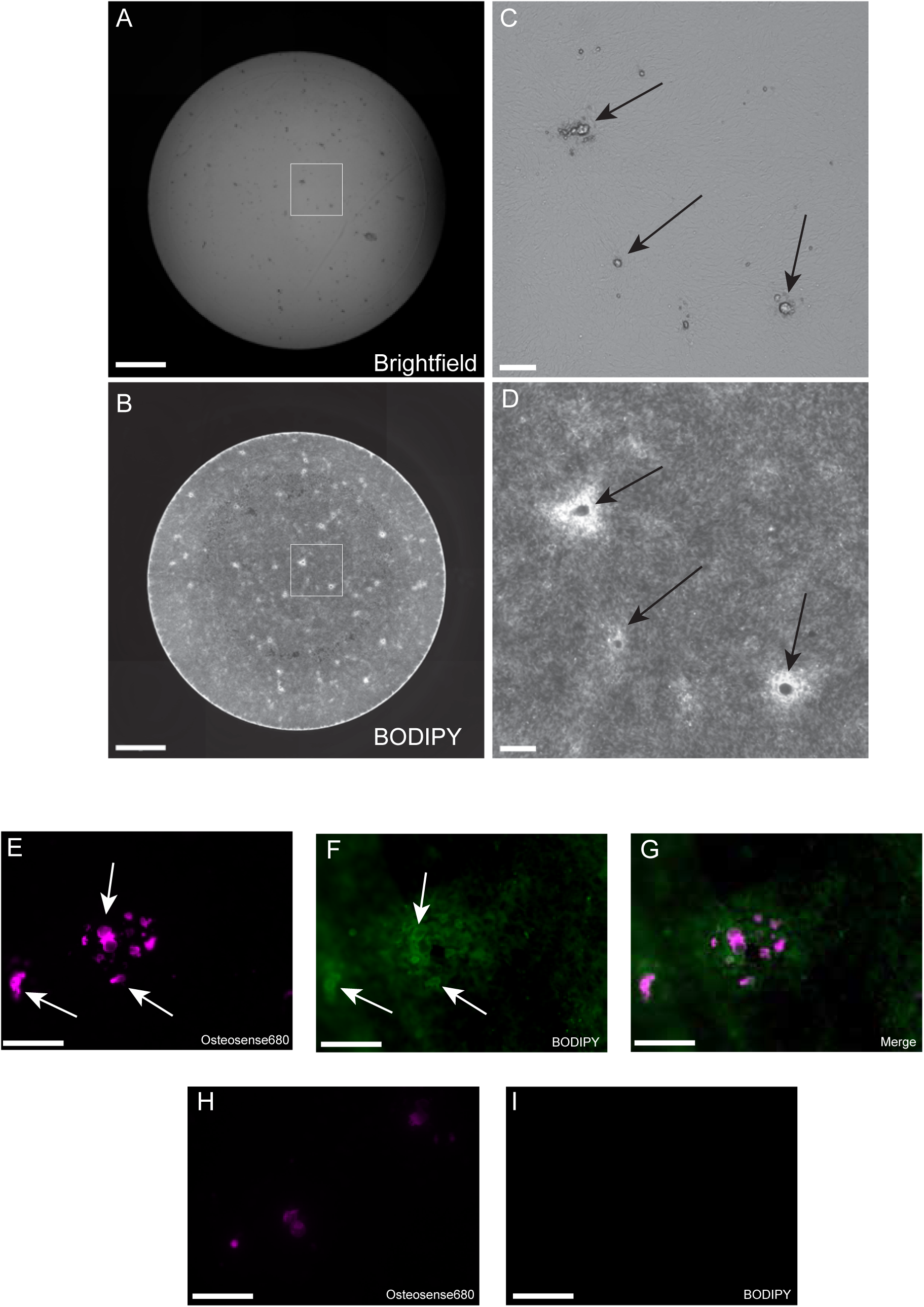
ARPE-19 derived HAP is associated with neutral lipid accumulation. (A-I) ARPE-19 cells were cultured in MEM-NIC + NaPAA conditions for 1 week followed by various forms of imaging. (A, C) HAP deposits in live-cell cultures were visualized by brightfield microscopy, with corresponding (B, D) BODIPY fluorescence showing neutral lipid-associated signal across the culture surface and surrounding discrete deposits. (E-I) ARPE-19-derived HAP deposits formed on transwells under the same conditions as in (A-D) showed punctate Osteosense680 labeling (E) and BODIPY-positive lipid-rich material (F). (G) Merged imaging of Osteosense680 and BODIPY. (H, I) No BODIPY control showes no signal in the GFP channel. Scale bars: A-B = 1000 µm; C-D = 100 µm; E-I, 100 µm. n=3 independent ARPE-19 staining experiments.

### ARPE-19 mineralization occurs without significant induction of osteogenic gene transcripts

To better understand the transcriptional changes associated with early-stage RPE-mediated HAP formation, bulk RNA sequencing was performed on ARPE-19 cells cultured in MEM-NIC in the presence or absence of NaPAA for 7 days. Differential gene expression analysis revealed distinct transcriptional changes between conditions, with 96 significantly upregulated and 71 downregulated genes identified following NaPAA treatment (Fig 4A, padj ≤ 0.05). Select transcripts that stood out based on their high fold change differential expression patterns or their low p-value were labeled on the volcano plot to illustrate high-confidence transcriptional change induced by osteogenic supplementation. Downregulated genes included naturetic peptide B (*NPPB)*, aryl hydrocarbon receptor R (*AHRR*), non-specific cytotoxic cell receptor protein 1 (*NCCRP1*), c-type lectin protein C-type lectin domain containing 19A *(CLEC19A*), and hypoxia-associated long coding RNA *MIR210HG* (Fig 4B, padj < 0.05). In contrast, upregulated genes including nuclear receptor subfamily 4 group A member 3 (*NR4A3)*, mitochondrial chaperone Heat shock 70 kDa protein 6 (*HSPA6),* cartilage glycoprotein chitinase-3 like-protein-1 (*CHI3L1*), and microsomal glutathione S-transferase 1 (*MGST1*) exhibited increased expression following NaPAA supplementation (Fig 4C, padj < 0.05). These labeled genes were not intended to represent a curated gene set, but rather transcripts that were visually notable within the overall differential expression profile. Absence of genes typically associated with mineralization such as *ENPP1, RUNX2, and TNAP/ALPL* may indicate that ARPE-19 mineralization differs from a classical osteogenic transcriptional response normally observed in, for example, vascular smooth muscle cells or osteoblasts.

**Figure 4.**
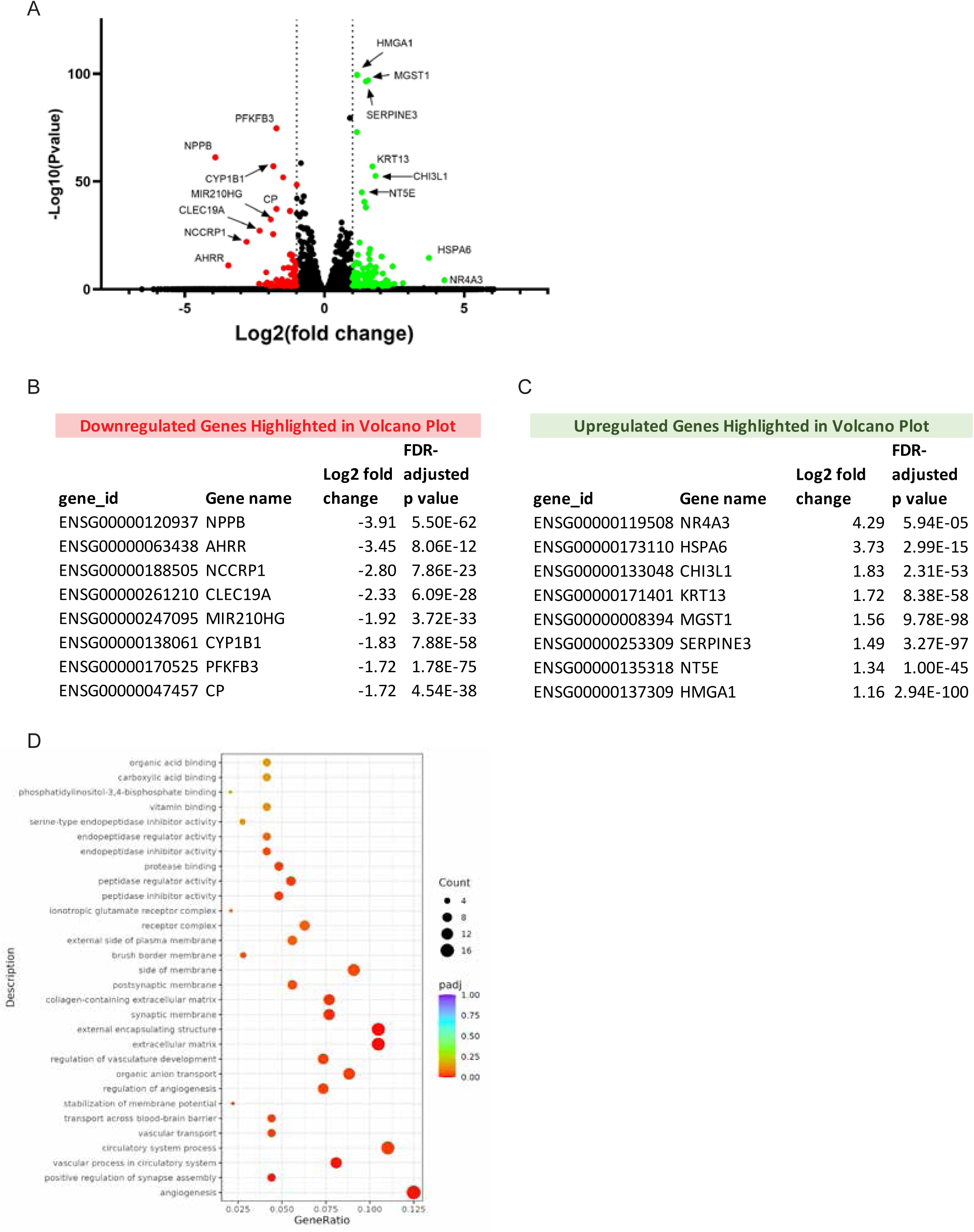
Bulk RNA sequencing reveals transcriptional changes associated with HAP formation in ARPE-19 cells. ARPE-19 cells were cultured in MEM-NIC in the presence or absence of NaPAA for one week prior to haresting for RNAseq. (A) Volcano plot showing differentially expressed genes in cells cultured in MEM-NIC + NaPAA vs. MEM-NIC alone. (B) Table showing the top significantly downregulated genes in MEM-NIC + NaPAA cultures compared with MEM-NIC alone. (C) Table showing the top significantly upregulated genes in MEM-NIC + NaPAA cultures compared with MEM-NIC alone n=2 biological replicates per condition. (D) Gene ontology enrichment analysis of differentially expressed genes shows an enrichment of angiogenesis, extracellular matrix, and circulatory system process. Dot size represents gene count, and color represents p-value (padj).

To identify biological pathways associated with ARPE-19 mineralization, gene ontology enrichment analysis was performed using differentially expressed genes. Enriched pathways included extracellular matrix organization, external encapsulating structure organization, angiogenesis, organic anion transport, and circulatory system processes (Fig 4D). Notably, gene ontology pathways typically associated with mineralization such as ossification and bone mineralization and were not indicated as enriched, suggesting that ARPE-19 mineralization under MEM-NIC + NaPAA conditions may occur through a non-canonical transcriptional program differing from bone or vasculature mineralization, in agreement with the notable absence of canonical mineralization genes indicated above. Overall, these results suggest that NaPAA supplementation induces a transcriptional response in ARPE-19 cells characterized by extracellular matrix-associated, inflammatory, and metabolic regulatory changes rather than activation of a canonical osteogenic-like gene expression program.

### Pyrophosphate and chelating agents significantly reduce ARPE-19 mineralization

Because ARPE-19 deposits exhibit chemical composition consistent with HAP, we sought to determine whether mineral formation could be disrupted by chemical agents. HAP formation depends on local calcium and phosphate availability, and mineral formation can be inhibited by chelating compounds that bind to and sequester metal ions such as calcium(de La Roij et al., 2026; Hentrich et al., 2017; Ramirez-Bommer et al., 2018). Pyrophosphate is a physiological inhibitor of mineralization and is found in the extracellular space where it functions to inhibit ectopic HAP formation in soft tissue and joints(Baptista et al., 2025; Liang et al., 2021; Orriss et al., 2016; Pomozi et al., 2017; Szeri et al., 2022). Pyrophosphate treatment reduced ARPE-19 mineralization in a dose-dependent manner, as measured by ARS staining and corresponding CPC extraction (Fig 5A–D). While 1 µM pyrophosphate produced what appeared to be a modest decrease in ARS signal, there was variability in the replicate data (Fig 5A, C). However, treatment with 5 µM pyrophosphate significantly decreased ARS staining, nearly prevented all ARS signal (Fig 5A, D). EDTA similarly inhibited mineralization, with both 100 µM and 500 µM EDTA significantly reducing CPC-extracted ARS signal compared with untreated controls (Fig 5E–H). Consistent with this quantification, post-ARS brightfield images showed a concentration-dependent reduction in ARS staining following EDTA treatment (Fig 5F–H). Sodium citrate also reduced ARPE-19 mineralization, however only a treatment 500 µM citrate significantly decreased ARS signal relative to untreated controls (Fig 5I–L). Representative brightfield images confirmed reduced ARS-positive staining following citrate treatment (Fig 5J–L). Together, these results show that ARPE-19 mineralization can be attenuated by pyrophosphate, EDTA, and sodium citrate, supporting the calcium phosphate-dependent nature of HAP formation in this model.

**Figure 5.**
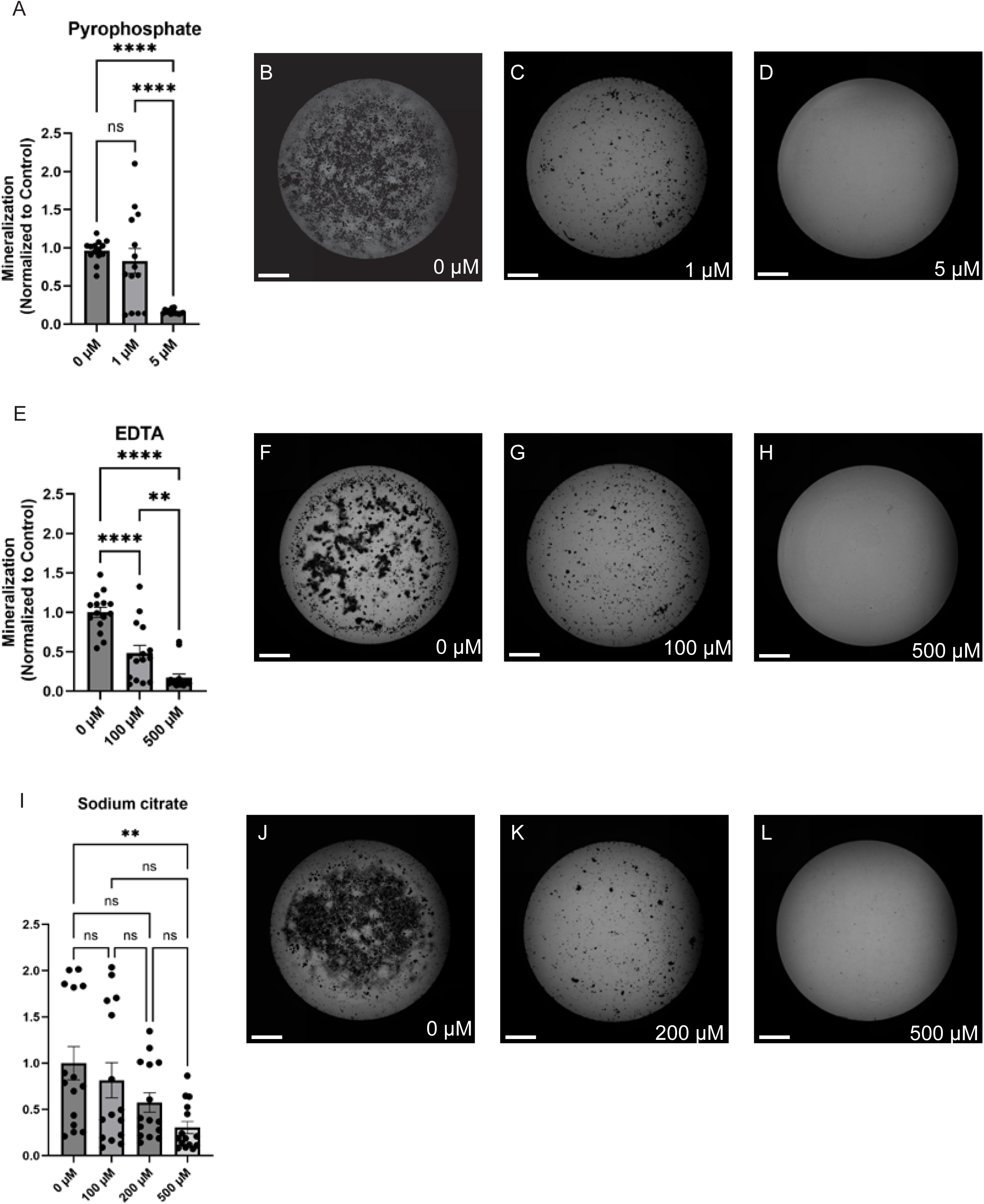
Pyrophosphate and chelating agents decrease ARPE-19 mineralization. ARPE-19 cells cultured under MEM-NIC + NaPAA were co-treated with with the indicated amount of pyrophosphate, EDTA, or sodium citrate to assess HAP formation for 14 days. Mineralization was quantified using cetylpyridinium chloride (CPC) extraction of ARS staining. (A–D) Pyrophosphate reduced ARS signal in a dose-dependent manner, with 5 µM significantly decreasing mineralization. (E–H) EDTA similarly reduced ARS signal significantly at both 100 µM and 500 µM, with strongest inhibition observed at 500 µM. (I–L) Sodium citrate significantly decreased ARS-based detecction of mineralization at 500 µM. Representative brightfield images of three independent experiments performed in quintuplicates are presented after ARS staining. Data in A, E, I are presented as mean ± SEM. Statistical significance was determined by one-way ANOVA. *p < 0.05, **p < 0.01, ***p < 0.001. Scale bars= 1000 µm. n= 3 independent trials on mineralized ARPE-19 cells.

## DISCUSSION

In this study, we established an inducible model of RPE-associated HAP mineralization using ARPE-19 cells. Importantly, the resulting cell culture-based deposits display characteristics reminiscent of those found in AMD-associated drusen. First, elemental analysis supports the relevance of the mineral produced in this model to that observed in human AMD tissue. ARPE-19-derived deposits demonstrated overlapping calcium and phosphorus enrichment by SEM-EDS and exhibited Ca/P ratios of approximately 1.50–1.52, compared with 1.73 in the human AMD-associated mineralization, and the idealized Ca/P ratio of 1.67 in HAP found in bone(Chokwattananuwat and Suttapreyasri, 2024; Lim et al., 2020; Mohring et al., 2023; Wang et al., 2010b). Although the Ca/P ratios found in deposits from mineralized ARPE-19 cells were not completely identical, their similar elemental composition supports the formation of calcium-phosphate-rich, HAP-like material under MEM-NIC + NaPAA conditions. Additional elements, including sodium, magnesium, aluminum, and chloride, were also detected within cultured deposits and human AMD-associated HAP. Some of these may originate from the culture environment or processing rather than representing intrinsic components of the mineral itself. Nevertheless, the similarity in calcium and phosphorus enrichment between ARPE-19 and human AMD-associated deposits provides important chemical validation that this system models aspects of the mineralization observed in disease. Moreover, while the majority of the deposits resemble spherules, some also resemble nodules associated with high-risk progression to advanced AMD(Tan et al., 2018).

Secondly, BODIPY-positive neutral lipid staining was enriched surrounding OsteoSense-positive deposits in mineralized cultures, reaffirming a proposed mechanism of drusen formation originating from previous studies of human AMD tissue. Previous work(Thompson et al., 2015) proposed that lipid droplets accumulating between the RPE and Bruch’s membrane may serve as nucleation sites for HAP deposition, which in turn could seed drusen formation, although this has not been directly demonstrated. Our findings would reinforce a spatial association between neutral lipid-rich material and HAP in an in vitro RPE system. Yet, longitudinal imaging will be required to determine whether lipid accumulation precedes HAP formation or instead occurs secondary to HAP deposition.

Our work provides the first set of bulk RNAseq transcriptional data originating from RPE cells mineralized under inducing conditions. At first glance, these observations suggest that the gene transcription profiles resulting from the mineralization process in these cells differs from classical osteogenic differentiation and vascular calcification, which typically involves upregulation of *ALPL, BMP2,* and *SOX9(Chen et al., 2022; Patel et al., 2019; Tian et al., 2025; Tyson et al., 2003)*. NaPAA supplementation under MEM-Nic media conditions produced transcriptional enrichment of pathways related to extracellular matrix organization, angiogenesis and circulatory processes. In contrast, regulators of osteoblast differentiation (*RUNX2 and SOX9)*, mineralization-associated genes (*ALPL and BMP2*), and extracellular-vesicle associated mineralization (*PHOSPHO1, SMPD3)* were not observed as differentially regulated. This observation is consistent with the inability of BGP to induce mineralization in our cultures, as BGP-dependent mineralization is characteristic of osteoblast and some vascular calcification models that relies on TNAP/ALPL activity. These findings raise the possibility that RPE-associated mineralization represents a form of pathological biomineralization that is mechanistically distinct from classical osteogenic calcification. In well-characterized models of vascular calcification, increased extracellular phosphate can promote osteogenic reprogramming of vascular smooth muscle cells, including induction of RUNX2 and increased activity of TNAP/ALPL, while simultaneously altering the extracellular environment in ways that favor calcium-phosphate nucleation and growth(Kapustin et al., 2015; Shanahan et al., 2011). TNAP is particularly important in this context because it hydrolyzes extracellular pyrophosphate, a potent endogenous inhibitor of HAP formation, thereby shifting the local balance toward mineral growth. These observations have established osteogenic differentiation and regulation of the extracellular mineralization environment as important, and sometimes overlapping, mechanisms of pathological calcification. Instead, the RPE may establish a local extracellular environment permissive for calcium-phosphate nucleation through changes in phosphate availability, extracellular matrix composition, cellular metabolism and secretion, lipid-rich or vesicular nucleation surfaces, or endogenous regulators of mineral growth. Such a mechanism would be consistent with the specialized biology of the RPE, which continuously regulates metabolite and ion transport(Skarphedinsdottir et al., 2020; Swarup et al., 2019; Yam et al., 2019), lipid processing(Lyssenko et al., 2018; Storti et al., 2017), extracellular matrix turnover(DiCesare et al., 2024; Eamegdool et al., 2020), and polarized secretion toward the sub-RPE compartment(Flores-Bellver et al., 2021; Lyssenko et al., 2018). The association between neutral lipid-rich material and HAP provides one potential example of how this extracellular environment may contribute to mineral formation. Although our experiments do not establish whether lipid accumulation, extracellular matrix remodeling, or other RPE-derived factors initiate mineral nucleation, they support a model in which the extracellular environment generated by the RPE contributes directly to HAP formation.

The extent of ARPE-19 mineralization could also be decreased by co-treatment using chelating agents. Pyrophosphate, EDTA, and sodium citrate each significantly reduced HAP formation when administered in conjunction with osteogenic supplementation, supporting the calcium-phosphate-dependent nature of the deposits. Although these agents act largely through physicochemical mechanisms and do not identify biological regulators of RPE mineralization, they provide proof of principle that that the model can be used to test interventions which modify deposit formation. These experiments also demonstrate that HAP formation can be experimentally modulated in this system. However, these treatments were only effective when present <u>during</u> mineral formation, whereas introduction after established HAP deposition did not result in a significant reduction in deposit size or number (data not shown), suggesting that these treatments primarily interfere with mineral initiation or growth rather than reversing established mineralization. This resistance, however, should not be evidence that established HAP deposits are completely irreversible. HAP is stable and poorly soluble under physiological conditions(Lacruz et al., 2017; Sakai et al., 2016; Wang et al., 2010b), and its resorption in bone requires osteoclast-mediated acidification and remodeling(Coury et al., 2013; Kartner et al., 2010; Riihonen et al., 2010; Supanchart et al., 2014). However, in some clinical cases of AMD, refractile or calcified drusen have been reported to regress without evidence of retinal atrophy(Liu et al., 2022; Oishi et al., 2017), suggesting that HAP deposits can undergo regression *in vivo* within the sub-RPE environment. The mechanisms underlying the cases of HAP regression in the retina, however, remain unclear. Together with our findings that chelating agents were only effective during mineral formation, these observations suggest that limiting HAP growth may be more tractable than reversing established deposits. For AMD, this raises the possibility that identifying smaller HAP deposits before extensive pathology occurs may provide a more effective window for intervention. At the same time, the spontaneous regression of some calcified drusen indicates that HAP deposits are not necessarily irreversible once formed and may remain subject to remodeling within the sub-RPE environment.

Several future directions emerge from this model. One priority will be determining the temporal sequence of neutral lipid accumulation and HAP deposition. Longitudinal imaging using BODIPY and OsteoSense, together with continuous live-cell imaging could define when individual deposits first emerge and whether lipid-rich structures precede mineral nucleation. Importantly, the inducible nature of this model provides an opportunity for high throughput screening of small molecules that either promote or inhibit mineralization. Unlike existing RPE models that require extended culture periods for mineral formation, this system offers a defined experimental timeframe suitable for screening large, small-molecule libraries. Such screens could identify modulators that either inhibit or enhance HAP formation and through subsequent target validation, reveal biological pathways that regulate RPE-associated mineralization. Candidate proteins within these pathways could then be tagged with HiBiT, enabling quantitative assessment of their endogenous protein abundance, responses to treatments, and HAP binding(DiCesare et al., 2024; Hulleman et al., 2025; Lankford and Hulleman, 2024). Together, this approach could help identify unrecognized pathways that regulate RPE-specific mineralization. Finally, convincingly reproducing the mineralization phenotype in primary human RPE, stem cell-derived RPE, or RPE progenitor systems will be important for determining whether the mechanisms identified in ARPE-19 cells extend to more physiologically representative models.

In summary, the major contribution of this study is the establishment of a controllable RPE-based system that generates HAP-like deposits with several features relevant to AMD-associated mineralization. The model permits mineral formation to be initiated within a defined experimental window, enabling the processes governing HAP initiation and progression to be studied separately. It also provides a platform for investigating how RPE-derived factors, neutral lipids, extracellular matrix changes, and mineral-regulating pathways contribute to HAP formation. Although this system does not fully recreate the aged sub-RPE environment (absence of immune or infiltrating cells), its experimental tractability provides an opportunity to dissect mechanisms that are difficult to resolve from human tissue alone and may ultimately clarify how RPE-associated mineralization contributes to drusen formation and AMD progression.

## ACKNOWLEDGEMENTS

We thank Nicholas Seaton, PhD, of the Characterization Facility at the University of Minnesota for his assistance with SEM-EDS instrument operation and data acquisition. This work was supported by a Training Program in Translational Vision Sciences grant (T32-EY025187 to JY), and R01-EY027785 (JDH). JDH is the Larson Endowed Chair for Macular Degeneration at the University of Minnesota and his laboratory is supported by the Helen Hessing Lindsay Family Foundation.

