## Supplemental Methods for "An Inducible Model of RPE-Associated Hydroxyapatite Mineralization with Relevance to Drusen Formation in Age-Related Macular Degeneration"

Seahorse assay. To measure mitochondria respiratory capacity, ARPE-19 cells were seeded on a Seahorse XFe96 (Agilent Technologies, Santa Clara, CA) plate at a density of 166,000 cells/cm^2^ with 10 µg/cm^2^ laminin coating (rhLaminin-521, Gibco). After 24 h, the media was changed to MEM-NIC or DMEM/F12 in the presence or absence of NaPAA and cultured for one week. On the day of the assay, cells were washed three times with Seahorse assay medium supplemented with 8 mM glucose, 2 mM sodium pyruvate, and 2 mM glutamine. Cells were then incubated in assay medium for 1 h at 37 °C in a non-CO₂ incubator. Prior to the start of the assay, mitochondrial stress test compounds were prepared to achieve final well concentrations of 2 µM oligomycin, 0.5 µM FCCP, and 0.5 µM rotenone/antimycin A (Agilent technologies). Seahorse data were analyzed using Seahorse Analytics software (Agilent Technologies). Statistical analyses and data visualization were performed using GraphPad Prism. Differences between experimental groups were assessed by one-way ANOVA followed by a multiple-comparisons test.

**FIGURE LEGENDS**

Supplementary Figure 1. Media composition of MEM-NIC and seeding density do not influence ARPE-19 mineralization. (A–E) ARS staining of ARPE-19 cells cultured in MEM-based medium containing different combinations of supplements, all with NaPAA for 14 days: (A) complete MEM-NIC media ; (B) MEM-NIC media without NIC, N1, (C) MEM-NIC media without NIC, N1, THT, (D) MEM-NIC media without NIC, THT, NEAA, (E) MEM base media without any additional supplements. Mineral deposition remained evident following removal of individual or multiple supplements. (F–H) Representative ARS staining of ARPE-19 cells seeded at different densities after 14 days: (F) 53,120 cells/well, (G) 6,640 cells/well, and (H) 3,320 cells/well. (I-J) OsteoSense680 staining was used to assess the presence of HAP deposits in MEM-NIC NaPAA-treated cultures in the presence or absence of ARPE-19 cells. (I) OsteoSense680-positive mineral deposits were observed in wells containing ARPE-19 cells, but not in the absence of cells (J). Scale bars A-H 1000 µm. n=3 independent experiments for MEM-NIC components and confluency tests. n=2 independent experiments for no cell osteosense staining.

Supplementary Figure 2. ARPE-19 mineralization occurs without changes in mitochondrial respiration. (A-E) Mitochondrial respiration was assessed using the Seahorse mitochondrial stress test in ARPE-19 cells cultured in DMEM/F12 or MEM-NIC in the presence or absence of osteogenic supplementation for 7 days. (A) Oxygen consumption rate (OCR) profiles following sequential treatment with mitochondrial stress test compounds. Quantification of basal OCR (B), ATP-dependent OCR (C), maximal respiratory capacity (D), and spare respiratory capacity (E). OCR values were normalized to Hoechst signal. Data are presented as mean ± SEM. Statistical significance was determined by one-way ANOVA. *p < 0.05, **p < 0.01, ***p < 0.001. n=4 independent experiments for Seahorse mitochondiral OXPHOS tests.
