## Supplementary figures and images for "An Inducible Model of RPE-Associated Hydroxyapatite Mineralization with Relevance to Drusen Formation in Age-Related Macular Degeneration"

### Supplemental Figures

#### Supplementary Figure 1

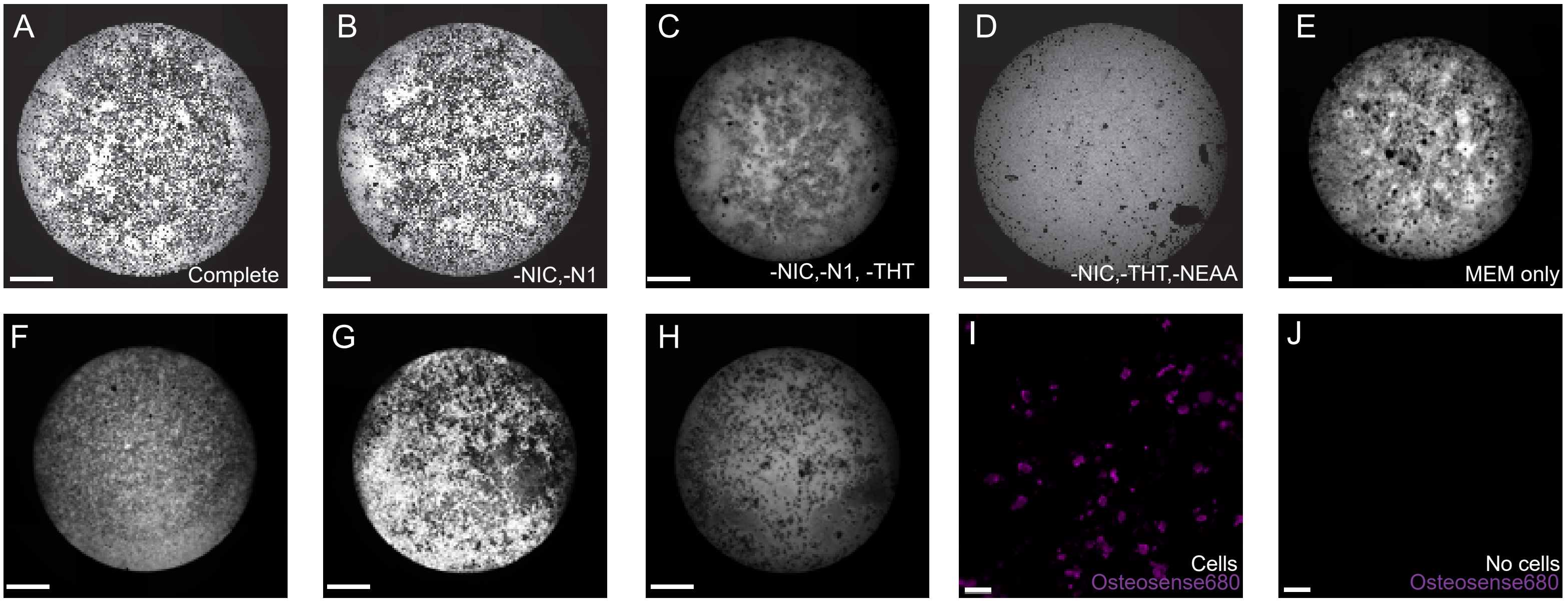

#### Supplementary Figure 2

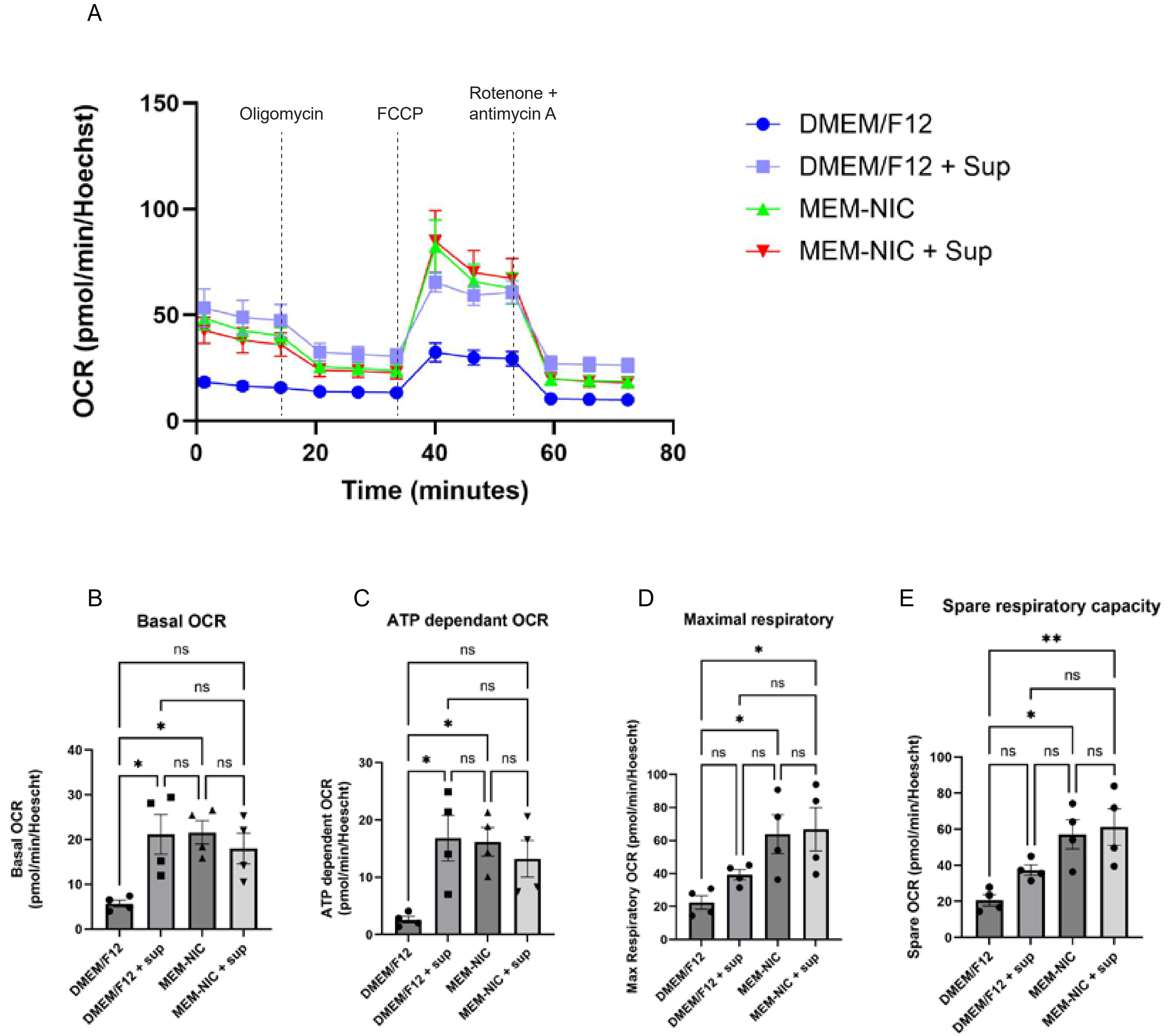
